# A common genetic architecture enables the lossy compression of large CRISPR libraries

**DOI:** 10.1101/2020.12.18.423506

**Authors:** Boyang Zhao, Yiyun Rao, Luke Gilbert, Justin Pritchard

## Abstract

There are thousands of ubiquitously expressed mammalian genes, yet a genetic knockout can be lethal to one cell, and harmless to another. This context specificity confounds our understanding of genetics and cell biology. 2 large collections of pooled CRISPR screens offer an exciting opportunity to explore cell specificity. One explanation, synthetic lethality, occurs when a single “private” mutation creates a unique genetic dependency. However, by fitting thousands of machine learning models across millions of omic and CRISPR features, we discovered a “public” genetic architecture that is common across cell lines and explains more context specificity than synthetic lethality. This common architecture is built on CRISPR loss-of-function phenotypes that are surprisingly predictive of other loss-of-function phenotypes. Using these insights and inspired by the *in silico* lossy compression of images, we use machine learning to identify small “lossy compression” sets of *in vitro* CRISPR constructs where reduced measurements produce genome-scale loss-of-function predictions.

## Introduction

Large-scale transcriptomics atlases have identified thousands of ubiquitously expressed mammalian genes^1^. Single-cell transcriptomics can rapidly enumerate the differential expression of the genome across a tissue^2,3^. Yet this unprecedented resolution in transcriptomics fails to tell us how genes function in different cell types. Importantly, while transcriptomics easily scales to measure thousands of transcripts in a single cell, functional genomics can only measure the phenotypic effect of a single genotype in a single cell. The knockout of a single gene can cause lethality in one cell and no discernible phenotype in another^4^. This diversity is even known to exist across KRAS mutant cell lines. Some G12X mutated cells are sensitive to loss of KRAS while others aren’t^5^. These phenotypic differences constitute a poorly understood, but well recognized phenomenon called context specificity that is important in cell biology, genetics, and medicine^6^. Context specific observations are a classic problem in mammalian cell biology and genetics that dates to the origins of the field ^7^. Understanding them aids our understanding of basic biological and genomic processes.

However, understanding cell specificity means contending with the fact that mammalian cell lines span an incredible diversity of potentially relevant contexts that include; nucleotide mutations, chromosomal abnormalities, copy number alterations, transcriptional profiles and tissues of origin^8^. Together, these contexts drive the richness and the confusion of mammalian cell biology/genetics. CRISPR screening efforts by the Sanger and Broad institutes have performed genome wide loss-of-function studies in nearly 1000 unique mammalian cell lines^4,9^. These studies have identified extensive phenotypic diversity (as measured by differences in CERES scores) in many genes^9,10^. Synthetic lethality has been the dominant paradigm that has been used to explore cell type specific sensitivity to loss-of-function in mammalian cancer cell lines. Classic synthetic lethality in cancer occurs when a specific somatic mutation confers sensitivity to genetic or chemical genetic loss-of-function^11^. This is best exemplified by the loss-of-function mutations in BRCA1/2 that predict exquisite sensitivity to the loss of PARP1/2. This relationship has led to the clinical application of PARP inhibitors^12^. Beyond this paradigmatic example, a number of interesting versions of synthetic lethality have been suggested, including; dosage lethality, paralog lethality, and lineage specific sensitivities^12–16,16,17^. All of these have been useful ideas to investigate the origins of cell type specificity and to create potential biomarkers for cancer therapy. However, the number of new and exciting synthetic lethality relationships discovered appears to be smaller than the phenotypic diversity observed in large scale cell line studies^6^, indicating that there is room for improvement in current association datasets.

One likely additional explanation for context specificity is that some molecular markers remain unmeasured. For instance, an unmeasured covariate is a potential explanation for any selective essential phenotype without a clear explanation. A second explanation is a lack of statistical power to identify rare variants or variants with modest effect sizes across large and heterogeneous groups of cells. Both these explanations can be tested and addressed by examining more data across more cell lines, and we applaud the ongoing efforts to do this. But, beyond these typical explanations, we posit that static genomic features of unperturbed cell lines may not be sufficient to predict context specificity. CRISPR knockouts create dynamic measurements of cellular responses to gene loss. This dynamic information may be useful to understand context specificity.

In this study, we explore the origins of cell type specificity as a question of basic mammalian cell biology and genetics. To do this, we take a data-driven approach that focuses on predicting cell type specificity with machine learning models. We carefully build thousands of machine-learning models that incorporated the effects of millions of CRISPR knockout phenotypes in addition to mutations, copy number, lineage, and RNA-seq to predict cell type specific phenotypes. Our analysis revealed that the best models of cell type specific CRISPR loss-of-function phenotypes were composed of other CRISPR loss-of-function phenotypes in a pooled library. Thus, “CRISPR predicts CRISPR” explains more context specificity than previous models. Furthermore, predictive CRISPR features fall into highly clustered and cross predictive subnetworks. When combined, these subnetworks constitute a “common genetic architecture”. While this architecture could be used for gene functional discovery, they also enable a powerful new approach for functional genomics. Inspired by the ideas behind data compression, we propose an approach for dramatically compressing genome-scale CRISPR functional genomics experiments. In lossy compression, orders of magnitude reductions in file sizes are exchanged for some acceptable tradeoff in data quality. Instead of *in silico* data, “CRISPR predicts CRISPR” models can compress *in vitro* CRISPR library composition. They identify reduced sets of CRISPR constructs that can predict the loss-of-function effects of unmeasured genes at tunable scales.

Chemogenomic screens bottleneck libraries by 10-fold at a 90% inhibition of cell viability and raise screening coverage requirements for genome scale screens to as many as 1 billion cells, a challenging bar. Pairwise genetic epistasis maps require *n^2^/2* unique constructs^18^. A genome-wide pairwise interaction map that uses the common coverage requirements of even 3 sgRNAs per gene and 500 cells/sgRNA and 500 reads/sgRNA creates impossibly large screening population in excess of 10^10^ cells. Thus, our lossy sets create an opportunity to extract genome-level information from smaller and more tractable libraries that will have applications in experiments that are impossible to scale to genome wide libraries. In doing so, we show it is possible to break the fundamental barrier of functional genomics; that only 1 genotype to phenotype relationship can be measured per cell in a pooled screen.

## Methods

### Overview of code repository

A fully documented git repository with all source codes and notebooks can be accessed at the Pritchard Lab at PSU GitHub page https://github.com/pritchardlabatpsu/cnp_dev. The source codes for running the entire pipeline have provided as python package, with all library dependencies defined for complete reproducibility – including the virtual environment used. All original data are already publicly available on the DepMap portal and upon download can be placed in the data folders as described in the documentation. The repo also contains codes for the generation of all figures/tables presented in this manuscript.

### Datasets

Cancer Dependency Map datasets 19Q3 and 19Q4 were retrieved from DepMap portal^19^. This included the Broad and Sanger gene effects (CERES scores), CCLE expression, CCLE gene copy number, CCLE mutations, and lineage (from sample info file). The classifications of nonessentials and common essentials were also retrieved for the same release periods.

### Data preprocessing

The data was pre-processed and later models were built using python. Several steps were performed to preprocess the raw data. Copy number missing value was replaced with zero. Cell lines with any missing CERES values were dropped. Mutations were grouped into damaging (with variant annotation labeled as damaging), hotspot non-damaging (with variant annotation labeled as not damaging and is either a COSMIC or TCGA hotspot), or other (with variant annotation labeled as other conserving/non-conserving).

Feature pruning was performed to remove non-informative variables. This included removal of features with only constant values and categorical features where if a category value was supported only by less than or equal to 10 samples. Non-expressed genes (TPM<1) were also removed. Data were normalized (z-scored) for RNA-seq and copy number values.

### Feature selection and model building

The model building was based on an iterative feature selection process. First, the dataset was constructed using CERES (excluding the target gene), RNA-seq, copy number, mutations, and lineage as features, with the goal of predicting the CERES of a target gene. The target genes list was a set of 583 genes with highly variable phenotype based on CERES standard deviation >0.25 and range >0.6. For lossy gene set compression inference (described below), the target genes list was the entire genomic set of 18,333 genes. The data was split 85% and 15% into train and test sets, respectively. The full training dataset was fit using random forest regressor with 1000 trees, maximum depth of 15 per tree, minimum of 5 samples required per leaf node, and maximum number of features as log2 of the total number of features. The top quartile of most important features were kept and used to refit a new random forest model. This process was repeated three times. The remaining feature set was further refined for significant features using the Boruta feature selection method^20^. The resulting features fit using random forest constituted the reduced model. For the purposes of analyses, the top ten most important features were selected and the resulting model constituted the top 10 only features model.

In addition to this iterative feature selection and boruta selection with random forest, other modeling approaches were performed for comparison. This included linear regression, elastic net (with alpha of 0.1), and random forest on the full train dataset followed by taking the top quartile of important features for building the reduced model. The top ten most important features were then taken for follow-up analyses.

### Model evaluation

The models were evaluated based on R2 values and were calculated for the full, reduced, and top 10 only features models. While R2 values provide a sense of model fit, it does not describe model significance. To this end, a recall metric was also calculated to take into account the correlation between predicted versus actual CERES relative to a reference null distribution. More specifically, recall was defined, similar to that in Subramanian et al, as the fraction of null Spearman correlation values that was lower than the Spearman correlation of the given model for the target gene of interest. The null correlation value was calculated as the Spearman correlation between the actual CERES of a randomly drawn gene and the predicted CERES of the target gene. This procedure was repeated 1000 times to generate the null distribution, per target gene. For the analysis set, we focused on predictable models with a recall >0.95 and R2 >0.1 of the top 10 only features models. In addition, a concordance score was calculated per model as the fraction of all predicted CERES values where the actual and predicted were both below or above −0.6 (a cut-off used for essentiality).

### Gene set analysis

For the examination of the relations between source and target genes in the models, several gene sets were used. Panther gene sets were downloaded from Enrichr libraries^21^. Paralog genelist were retrieved from Ensembl gene trees using *biomaRt* in R^22,23^. The HGNC symbols of the most diverse genes were used as filters and the queries were submitted to GRCh37 Homo Sapiens Ensembl Genes *biomaRt* database. Gene set enrichments were performed using *gprofiler2* in R with default parameters and a p-value significance threshold of 0.05^24^.

### Network analyses

Networks were derived from the model results by linking all related source and target genes using *networkx* in python. Communities were extracted from this network using the Louvain method. Network communities were visualized and network statistics were extracted using Cytoscape v3.7.2 ^25^. Power analysis was performed on nodes with undirected degree of at least 2. The number of nodes vs degree was fit to a power-law function y=ax^b^.

### Lossy gene sets

Lossy sets were derived as centroids in an iterative k-means-based tight-clustering procedure using the *tightClust* package in R^26^. The procedure was run to derive L25, L75, L100, L200, and L300 lossy sets, which were then used for the saturation analyses. For compression-based inference, the input dataset consisted of the CERES of these lossy gene sets, as opposed to all genomic features.

### Statistical analyses

The enrichment of contribution of CERES as features were statistically tested using Chi-squared test. The statistical significance for node clustering coefficient and average number of neighbors between functional only and functional+genomic networks was assessed using two-sided t-test. All statistical tests were performed in python.

## Results

### Cell type specific loss-of-function phenotypes are predictable with machine learning

Recent efforts at the Broad and Sanger institutes have created an unprecedented resource to investigate the origins of context dependence across mammalian cell lines. Context specific behavior in CRISPR loss-of-function phenotypes are defined across cell lines using CERES scores^9^. Briefly, a CERES score measures the size of the fitness difference that is observed in a pooled screen from a single mammalian cell line. CERES scores are calculated for every gene in the library and they account for multiple confounding effects that bias the direct measurements of individual sgRNA enrichment and depletion. Negative CERES scores denote net gene depletion and positive scores denote enrichment across sgRNAs. All CERES scores for a gene across all cell lines provide a high resolution measurement of cell type specificity.

In considering how to model cell type specific phenotypes from CERES, genes could follow 2 distinct models of cell specificity. For example, in one widely used model, CERES scores are thought to belong to 2 distinct probability distributions that are simply “essential” or “not essential” for growth in an individual cell line. This first model suggests that individual cell lines can be assigned 1 of these 2 classes for any gene. However, CERES scores in common essential genes can harbor large variation across cell lines^19,27–29^. Moreover, some genes that are labeled “common essential”, “selective essential”, and “non-essential” have large ranges of continuous variation in CERES scores that appear to be biologically meaningful **(Figure 1A,B)**. Thus, an alternative explanation is that CERES scores are continuous outputs of a single cell type specific function across cell lines that represent the degree to which gene loss-of-function changes cell growth rates^30^.

**Figure 1.**
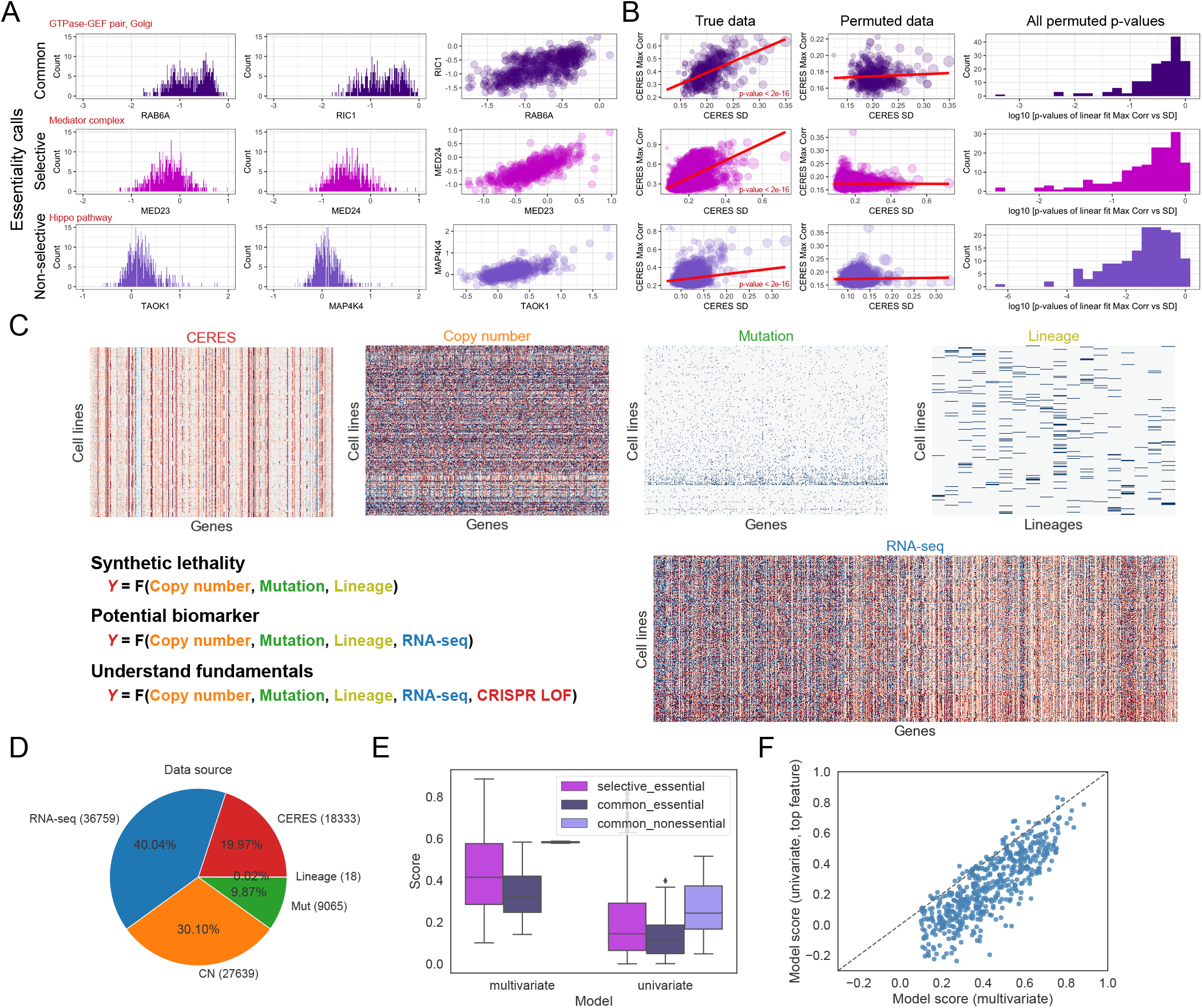
Multivariate machine learning models predict context specificity. (A) Examining the largest and most cell type specific genes in the DepMap data set identified genes for whom CERES scores were highly correlated with a biologically related gene in the same pathway or complex. This was true in common, selective, and non-essential DepMap genes. This suggested that continuous variation in CERES scores can have biological meaning (B) Scatter plots (left, middle) have 18333 unique genes as data points. The standard deviation of CERES score for each gene is plotted on the x-axis, and the point size is proportional to the range of CERES scores across all cell lines. The most cell type specific genes tend to have the strongest correlations with CERES scores in other genes. This is measured via a linear model (red line), and the p-value of this model is <2e-16 in the true data (left) for common, selective, and non-essential genes. To assess significance of the relationship between standard deviation within a gene and biological correlations with the CERES scores of another gene, 1 example of a permuted CERES dataset and its linear model is shown in the middle. 163 permutations of the CERES scores and the p-values from their respective linear models are shown in the histogram to the far right. Permuted p-values rarely drop below 0.001 and are never below 10^−6^ while the real data had a p-value of <2e-16. The more context specific a gene is, the more likely it is to be highly correlated to a second gene. Correlation with a second gene is an indicator that CERES score variation has biological meaning. (C) Heatmaps of datasets available for the prediction of context specificity from the 19Q3 DepMap CERES scores, copy number, RNA-seq, mutation, and lineage. The colors were scaled to be between −1 and 1. We have focused on the use of CERES scores in addition to traditional genomic features as potentially useful for predicting context specificity in loss of function phenotypes. (D) The distribution of number of features per data source. (E) After cross validation and validation of test set predictive power, models for each context specific gene are plotted as individual points. 1 data point is 1 gene. Multivariate models(x-axis) are more predictive than univariate models(y-axis), across all gene essentiality classes. (F) Comparisons in model scores between the multivariate versus univariate models across different essentiality classes. Boxplots are broken down by gene classification, score is based on performance during cross validation.

To be agnostic to these two models, we used standard deviation and range cutoffs across the CERES scores for all genes in all cell lines. We found that genes with more context specificity across cell lines (i.e. increasing standard deviation and range), tend to have highly correlated non-paralogous genes in the dataset **(Figure 1B)**. This relationship was not simply a function of increased variation because permutation tests failed to observe similar trends in the data. We visualized distributions and identified 583 genes out of 18,333 total genes using a cutoff (see methods) that identified the largest standard deviations and the most expansive ranges CERES scores across all cell lines. **(Figure S1-a-A)**. These highly context specific genes were enriched for functions in the cell cycle, metabolic processes and mitochondrial processing. (**Figure S1-b**).

There are multiple potentially predictive data types (and millions of data points) assayed in the DepMap project that could be used to infer the origins of context specific loss-of-function phenotypes **(Figure 1C,D)**. All of the datasets (mutation, RNA-seq, CNV, and lineage) that we examined from the CCLE contained a large dynamic range of measurements **(Figure S1-a-B)**. This suggested that variation in any/all of the data types could be useful for making context specific predictions of CRISPR loss-of-function phenotypes and inferring predictive features of interest.

Traditional definitions of synthetic lethality in cancer cell lines examine univariate genetic predictors that include copy number and mutation context **(Figure 1D,E)**. Beyond classic definitions of synthetic lethality, lineage and transcriptional data have also been used to mine large cancer datasets. Because we aim to understand the nature of context specificity and how predictable it is, we took a fundamentally different approach from recent impressive work that ranked potential univariate biomarkers and drug target pairs^4^. Compared to univariate predictors, our first question was whether multivariate machine learning models can explain larger amounts of context specificity. We also relaxed the requirement for translational utility by allowing a CERES score in one gene to be predicted by other CERES scores across the genome. We did this because we suspected that the collective probing of the loss-of-function dynamics of the entire genome through CRISPR could provide information on the state of the genetic network that is fundamentally different than baseline OMICS measurements in unperturbed cells.

With millions of measurements, we needed to build an extensive feature selection and model validation pipeline for our machine learning models to minimize overfitting while ensuring robustness and predictive power (**Figure S1-c**). Comparing across multiple machine learning methods, we found that an approach that employed iterative-feature selection combined with random forest regression was robust and superior to other approaches (**Figure S1-d and Fig S1-e**). We validated our models with additional test sets (from later DepMap releases) and we observed comparable results (**Fig S1-e A-C**).

With this high-quality pipeline, we built thousands of machine-learning models across millions of measurements. We found that multivariate models virtually always outperform univariate models when predicting diverse cell line specific phenotypes across our set of highly context specific genes **(Figure 1E,F)**. This suggests that multivariate context critically improves predictions of cell type specificity, and that classic synthetic lethality as a univariate paradigm to interpret context specificity can be greatly improved upon. We observed this for models that predicted cell line specificity as essential or not, and models that predicted cell line specificity as a continuous CERES score. However, are our multivariate models better because they add further genetic context to key synthetic lethal mutations? Or is the multivariate prediction accuracy due to something else?

### “CRISPR predicts CRISPR” rationalizes cell line specific responses to gene loss

To investigate why our multivariate machine learning models make better predictions than univariate models we first examined the types of data that contribute to highly predictive models of context specificity. A plurality of input features (40.0%) were RNA-seq transcripts in the input data set, but the top 10 features in our predictive multivariate models were overwhelmingly composed of CERES scores (73.4%) (p-value < 0.001) (**Figure 2A-B, and Figure S2-a A and Figure S2-b**).This suggested to us that perturbations of the genetic network (as provoked by Cas9, and measured by CERES scores) were better at predicting cell type specific phenotypes than conventional “omic” measurements from unperturbed cells.

**Figure 2.**
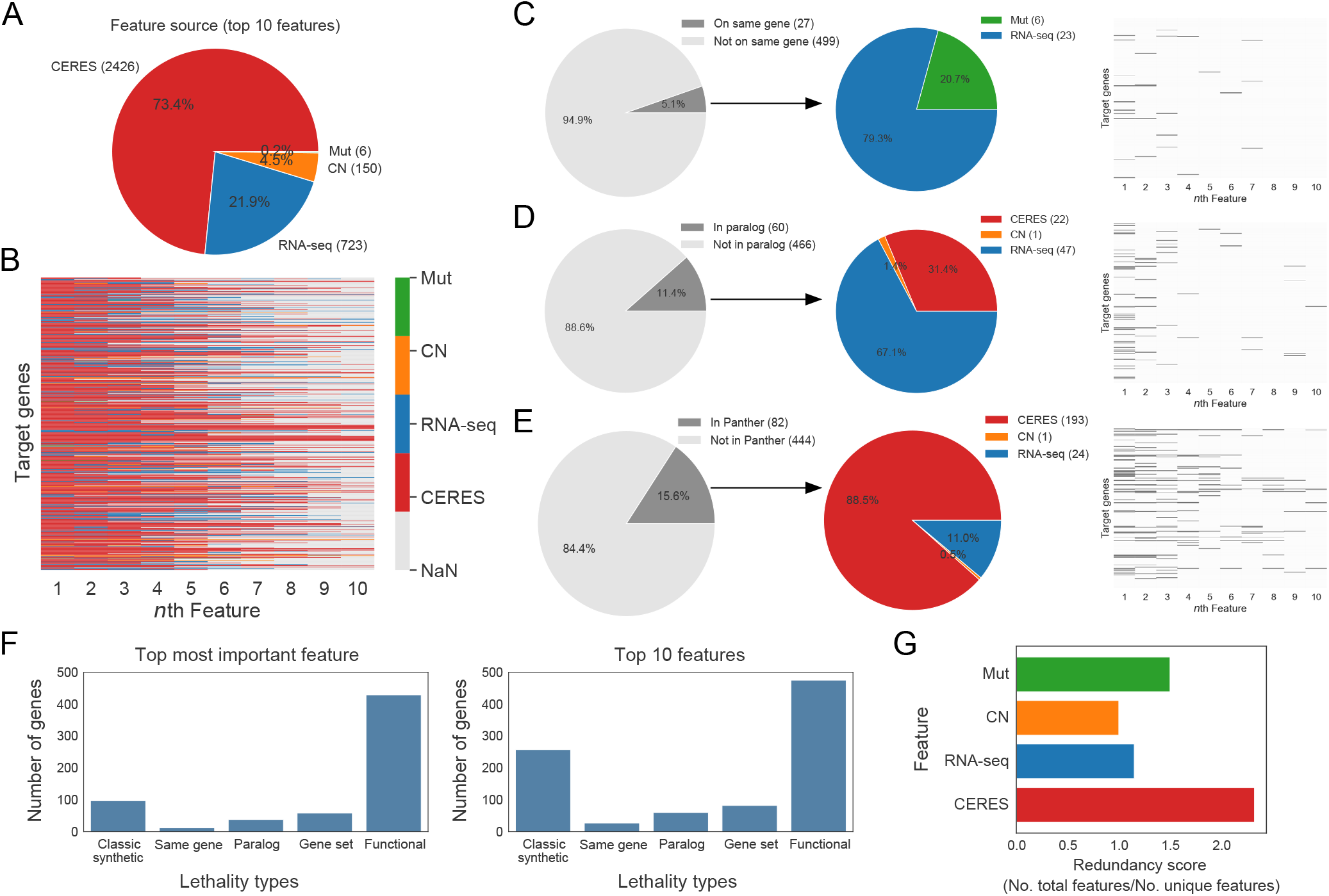
CRISPR CERES scores are the most prominent predictive features in multivariate models. (A) The distribution of the top 10 features per data source from the multivariate models in figure 1 are aggregated across all models. CERES scores are highly enriched as features in these multivariate models. (B) A heatmap view of all target genes for *n*th feature, colored by data source. (C-E) The relationship between the top 10 features for every cell type specific gene and whether they are from the same gene (C), paralogs (D), or in network gene set Panther (D), as visualized in the gray pie charts and heatmaps. Of the feature-target gene pairs within each group (dark grey), they are further broken by data source (colored pie charts). (F) Examining the top feature in every model, we classified which model of cell type specificity that feature belonged too. (G) Redundancy scores are measured as the ratio of the total number of features to total number of unique features, measured per data source (CERES, RNA-seq, mutation, CNA, lineage).

To quantify the degree of these differences and to understand them, we examined our multivariate models of context specific loss-of-function using existing literature models of context specificity. These alternate models include classic synthetic lethality (the genotype of cell x predicts sensitivity to LOF), dosage lethality (the expression level of a gene predicts its own sensitivity to knockout), paralog lethality (the loss of a compensating paralog confers sensitivity to a related gene), lineage lethality (where cell type or developmental context predict genetic sensitivity) and network context (where neighboring genetic nodes in a network diagram are connected using another database that adds signal to noise in context specific predictions)^4,14,31–33^. Amongst the top 10 features in our multivariate models, all of these existing models of context specificity were observed (**Figure 2C-F**). This is consistent with exciting recent papers where similar approaches have featured prominently^4,31,33^. Moreover, it re-emphasizes the importance of these relationships in a select group of context specific genes. However, despite our replication of previous paradigms, we found >4 times as many context specific CRISPR loss-of-function phenotypes were accurately predicted by CERES scores in another gene in the genome (**Figure 2F**). We term this finding “CRISPR predicts CRISPR”.

Therefore, prior paradigms do not capture as many context specific relationships as our newly discovered “CRISPR predicts CRISPR” approach. Importantly, this also represents a distinct model of how context specificity can be generated. In classic synthetic lethality, cell specificity is predicted by “private” mutations that are unique to a subset of cells. Synthetic lethality suggests that these private mutations cause a genetic network to function in a different way in 2 different cell types. In contrast, our CRISPR predicts CRISPR models utilize measurements of the function of predominantly wild type genes that are consistently expressed across cell lines, only 6.9% of genes with significant CERES scores have any alterations in the omic data in any cell line. The collective probing of these “public” nodes that are present across most, if not all, cells identifies a different way to explain context specificity. It implies that many genetic rules are shared across all cell lines and that cell line specificity can be driven by the degree of common pathway utilization. Importantly, this is true in genes that are labeled as “selective essential” and “common essential”.

To understand this common genetic architecture in more detail we also sought to understand the topology of the combined set of all multivariate models. In other words, we asked how many predictive features are shared between models. If multiple models share the same CRISPR features it would suggest that some genes might be especially good at predicting context specificity. It would also suggest that some screening information might be redundant. We divided the total number of predictive relationships by the number of unique predictors to get the average number of predictive relationships per feature. CRISPR features had an average of 2 relationship/feature, meaning that unlike mutation predictors, many CRISPR gene features were shared across multiple models (**Figure 2G**).

### Densely cross predictive CRISPR networks define a common genetic architecture in mammalian cells

Because of the dichotomy between the “private” features that predict synthetic lethality, and the “public” features in our CRISPR predicts CRISPR models, we provide a schematic that describes our hypothesis to account for these differences. In cancer models of synthetic lethality, a variety of possible models exist. These distinct relationships form distinct private functions *f_i_(x)* that predict cell type specific differences in CERES scores *Y_i_* **(Figure 3A, Left)**. The predictive features in these functions are rarely shared between models **(Figure 2G).** However, in a common genetic architecture these private functions exist in the background of a densely predictive common genetic architecture that ties many individual *Y_i_* predictions together, and many CERES features are shared between models **(Figure 3A, Right)**. This hypothesizes a different network structure. We asked whether this schematic was consistent with our models of the DepMap data by using the union of all models to build networks where genes are nodes, and predictors of cell type specific phenotypes are edges **(Figure 3B)**. When we compared the networks with and without CRISPR CERES scores we observed a striking dichotomy **(Figure 3C)** that matched our hypothesis from **Figure 3A**. CRISPR CERES predictions connect many private models into larger “public” subnetworks **(Figure 3C,D)**, this dramatically increases the number of nodes (genes) that exist in the network (i.e. It explains more context specificity), and these nodes have a higher mean number of neighbors 5.1 vs 1.7 (p-value = 2.0e-24), and a higher clustering coefficient 0.5 vs 0.04 (p-value = 1.7e-40) **(Figure 3E,F)**. These highly connected and densely clustered subnetworks identify the fabric of genetic interactions that are interwoven to collectively determine cell specificity across cell lines.

**Figure 3.**
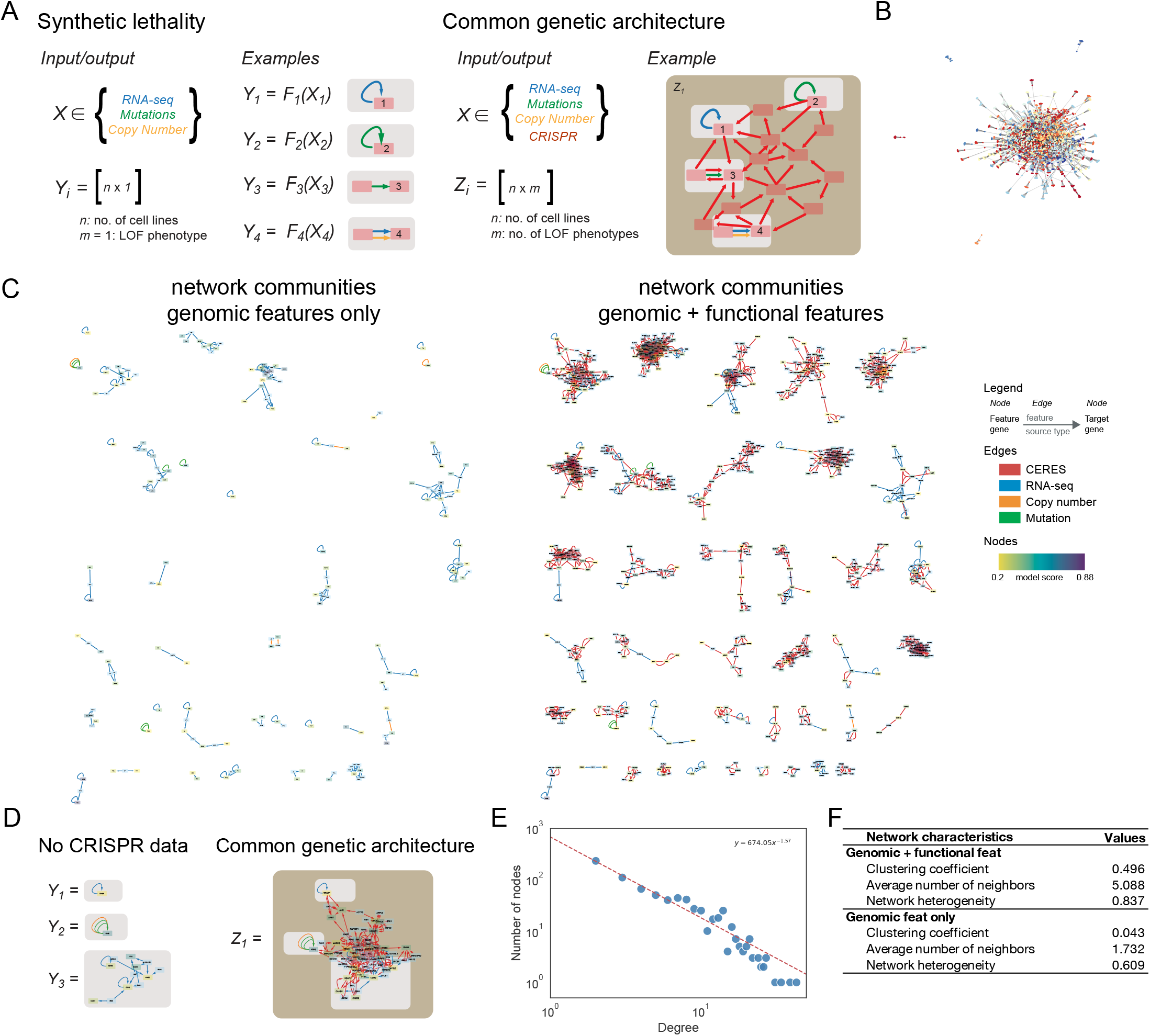
A common genetic architecture in mammalian cell lines. (A) Schematic illustrating difference between classic synthetic lethality and our *common genetic architecture*. Synthetic lethality consists of many individual *Yi* functions. These functions are cell type specific “private” models. Our proposed common genetic architecture is hypothesized to connect these private functions with shared CERES features. A common genetic architecture has redundant edges, and more nodes. More nodes suggest that more cell type specific phenotypes are predictable. (B) A network built from the aggregation of all multivariate models. Genes are represented as nodes and feature-target gene relations as edges. Colors represent distinct subnetwork communities that were identified by the Louvian method. (C) Network communities with (right) and without (left) nodes/edges involving functional CRISPR features. Edges are colored based upon the data source; and nodes are colored based on the model score (of top 10 feature model) of the corresponding gene as target. (D) Illustration, based on top left network community from (C), in terms of the individual *Y* and aggregated *Z* functions. (E) Number of nodes versus node degree based on the network genomic+functional features. The data is fit to a power law function. (F) Characteristics of networks with and without CERES features.

These networks obey the power law (**Figure 3E**), and highlight coherent biology (**Figure 3C, S3-a, Supplementary Data**). For example, **Figure S3-a-A** shows a Cyclin-CDK regulatory network that is entirely composed of genes that drive the cell cycle. **Figure S3-a-B** highlights mitochondrial respiration and the electron transport chain. Thus, our common genetic architecture identifies coherent biological subnetworks with highly cross predictive CERES scores that could be used to annotate the function of unannotated genes. But our most important hypothesis is that not all genes in a subnetwork would need to be measured in order to predict the CERES scores across an entire subnetwork.

### Identification of a common genetic architecture leads to the lossy compression of CRISPR libraries

Measuring sgRNA enrichment or depletion in pooled screens informs gene function by identifying when genes are required for viability^27,28,34^. However, these libraries can be too large and cumbersome for many types of biological studies. This is because CRISPR libraries require large amounts of coverage (4-10 sgRNAs/gene, 500 cells/sgRNA, and 500 reads/sgRNA) to achieve high quality measurements^18,35,36^. A “simple” genome-wide dropout screen in a mammalian cell line can require 40 to >100 million cells and a similar number of sequencing reads to get good measurements on nearly 20,000 genes. Thus, an approach that decreases the measurement requirements at the genome-scale would be useful for mammalian cell biology and genetics. This reduction creates a natural analogy to image compression. “CRISPR predicts CRISPR” models might be able to compress CRISPR experiments instead of their data. We aim to identify reduced sets of CRISPR constructs that could be screened at smaller scales to predict the loss-of-function effects of unmeasured genes at the genome-scale.

Image compression can be lossless or lossy. Lossless compression achieves modest reductions in file sizes but loses no information. Lossy compression can achieve orders of magnitude reductions in file size **(Figure 4A)**, but there are tunable reductions in image quality^37^. For our purposes, a dramatic reduction in library size with some information loss is preferable to lossless compression and a modest reduction in library sizes. This is because a fundamental barrier to the scalability of modern human functional genomics is library size ^18^. Achieving less than an order of magnitude reduction in library size will be useful, but it will fail to dramatically enhance experimental tractability in many experimental contexts. For instance, even a genome-wide library of perfect sgRNAs cannot decrease CRISPR library sizes to below ~20,000 unique constructs. Eliminating the need to measure redundant genes (and not redundant sgRNAs) is the only approach that has the potential to create libraries that are orders of magnitude smaller.

**Figure 4.**
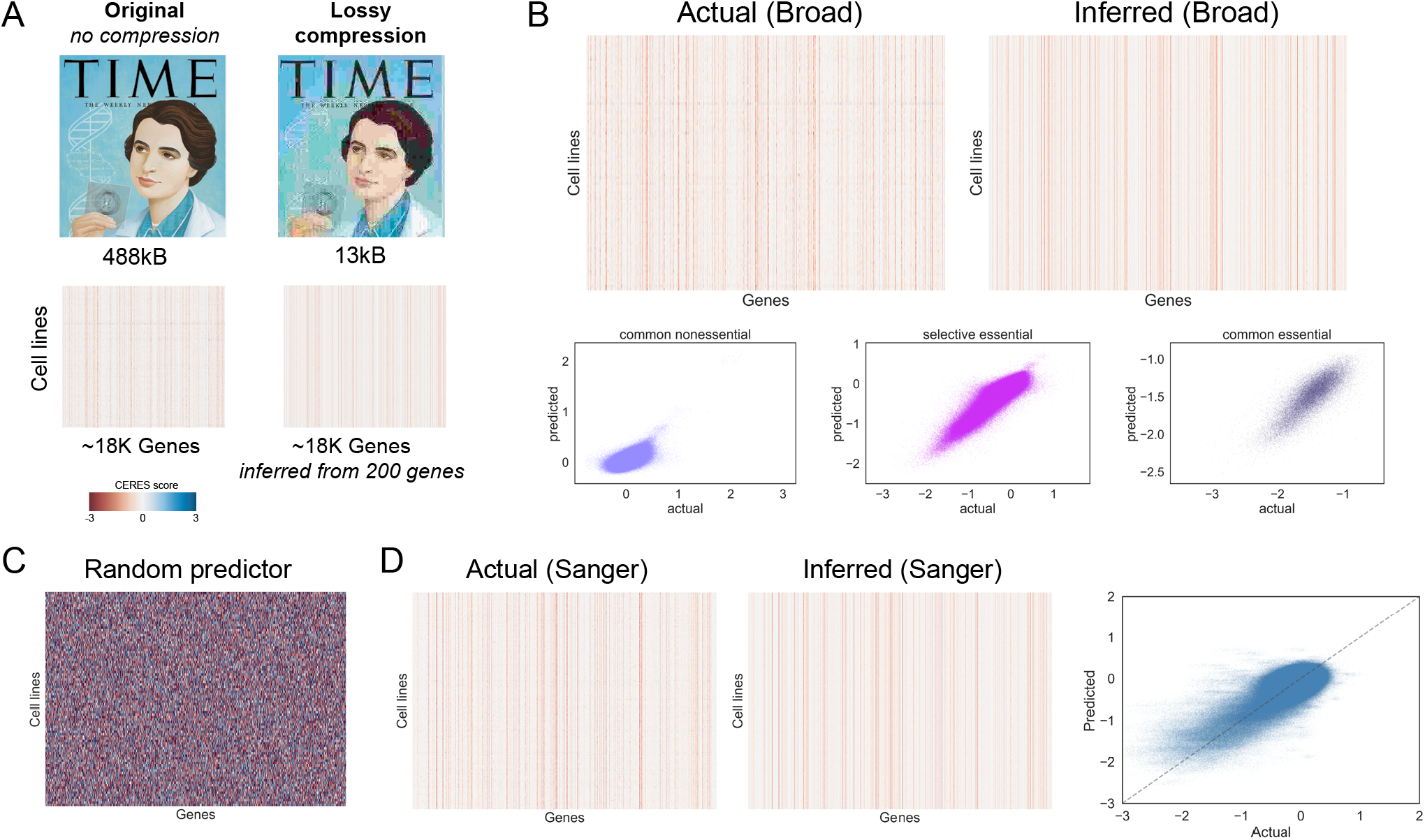
Lossy genomic compression. (A) Schematic of lossy compression in terms of image compression using JPEG (top) and similarity to genomic loss-of-function compression. (B) Genome-wide comparison between actual and inferred CERES scores on a held-out test set, shown as heatmaps and scatter plots per gene essentiality classes. Models built based on only 200 lossy genes with their CERES scores as features. (C) The same test set as in (B) but was inferred based on a random predictor. (D) Genome-wide comparison between actual and inferred CERES scores on a held-out test set from Sanger. Multivariate models were trained based on Broad data.

Inspired by lossy compression and previous landmark analyses in transcriptomics^38^, we hypothesized that the existence of a common genetic architecture could enable the tunable lossy compression of the genes that are targeted in CRISPR libraries. Here compression does not refer to the *in silico* data storage requirements, but to the *in vitro* number of genes that must be measured in a pooled screen to achieve a genome-scale “portrait” of gene function (**Figure 4A**).

To examine the possibility for lossy compression we decided to start with the entire genome, and not simply focus on the genes that were highly cell type specific. Our aim was to identify a highly informative set of genes, for whom experimental measurement can accurately predict unmeasured CERES at a genomic scale. Using a multi-step machine learning approach, we identified 25, 100, 200, and 300 gene sets whose CRISPR CERES scores were highly correlated with other genes in the 19Q3 Broad DepMap data. These potential compression sets were enriched for genes that are involved in metabolic processes, nuclear organization, organelle biosynthesis, and transcription factor activity **(Figure S4a-A-C)**. We then examined whether these small gene sets harbored predictive power across the genome. With models trained on 488 cell lines and then tested on 87 independent cell lines screened months later, we found that the predictive performance of lossy compression genes in a validation set began to saturate at 200 genes **(Figure S4b-A)**. Across the genome, predictions from a lossy 200 set closely resembled the true test set data (Pearson r = 0.92) (**Figure 4B and S4-b-B-D).** Importantly, the similarity between predicted and measured CERES scores existed across common essential, common nonessential, and selective essential genes. The predictions were dramatically better than models built from a random set of 200 genes (Pearson r = 0.0) (**Figure 4C)**. To build intuition, 2 representative examples of 2 gene specific models are available in **Figure S4b-D** and all 18,333 models can be reproducibly generated from source on GitHub. Importantly, some common essential and nonessential genes were predicted to have roughly the same phenotype in all cell lines. This is consistent with the interpretation that some essential genes are interpreted in a binary fashion by the cell. This can be seen in our use of qualitative comparisons of accuracy in addition to observed vs predicted plots.

Finally, recent work comparing the Broad dataset with the Sanger dataset has identified high cross-dataset correlations and key differences for many shared genes^10^. These projects had differences in screen duration, media composition and sgRNA identity. As an extraordinarily stringent test of our lossy 200 set, we examined the predictive performance of our lossy 200 gene set that was built on Broad data in a second test set from the Sanger institute **(Figure 4D)**. Consistent with a lossy approach, our lossy 200 set reproduced a genome scale portrait of CRISPR loss of function in the Sanger institute data that resembled performance in the DepMap test data (Pearson r = 0.78). This is despite the Sanger institute’s use of different sgRNAs, and different screening conditions that our lossy 200 gene set was not trained upon. This clearly suggests that small sets of 200 genes can provide reproducible genome-scale information.

## Discussion

What makes different cell types behave differently? This has been a fundamental question of genetics and cell biology since the first cells were cultured. It has guided classic work by Hayflick and Eagle^7,39^, as well as modern work in systems biology^40^. Recent systematic efforts to perform genome scale CRISPR screens across mammalian cell lines are a significant new tool to understand this challenging question. This is due to the exceptionally detailed and comprehensive genomic information that now exists in CRISPR screened mammalian cell lines across 2 independent institutes^4,8,9^. Our work represents the largest systematic effort to use these datasets to understand the basic biological origins of cell type specificity. Understanding this question advances cell biology, genetics and cancer biology. Previous work has focused on ranking translational hypotheses and comparing dataset measurement quality^4,10^. Surprisingly, well described and clinically important phenomena like synthetic lethality can make strong models, but they are relatively rare explanations for context specificity across all cell lines/genes. We term them “private models”, where a specific event in a subset of cell lines predicts a unique response to CRISPR mediated loss-of-function. Thus, private genetic mutations constitute a comparatively minor explanation for cell type specific phenotypes.

Instead of private models, the collective wisdom of all CRISPR loss of function perturbations across multiple genes constitutes a “public” model of cell type specificity where the elements of genetic logic are shared across cell lines. Cell type specificity is produced by the differential utilization of underlying genetic pathways that are common across cells. These public models are based upon the information encoded in the response to stimuli in widely expressed genes and not mutations in those genes. They can also be collectively assembled into a common genetic map that details the network architecture of cell type specific phenotypes across cells. Interestingly this common map resembles a decade-old concept in the signal transduction literature called “common effector processing”^40^. Common effector processing describes cell specific differences in caspase activation as a function of signal integration across a small set of widely expressed kinases and not the differential utilization of cell type restricted kinase pathways. Thus, our genetic findings converge with classic high-profile work on post translational modification networks. This convergence points to a broader theme in explanations of context specificity that transcend a single data type or phenotypic focus. This theme is that while discrete and qualitative differences between cells can drive cell type specific behaviors, it is often the quantitative degree to which ubiquitous pathways are utilized that determines context specificity. This is a continuous model of cell specificity in public genes, as opposed to the discrete model of synthetic lethality in private contexts. Our systematic analysis across the Broad and Sanger datasets lends support to this model of context specificity at a scale that dramatically exceeds prior work.

In network biology, previous work has focused on the most connected nodes. So called “Network Hubs” are often enriched in functional phenotypes and regulate many genes^41^. In our study, the most connected hubs contain the most redundant features in our predictive models. This dichotomous interpretation of hubs (critical vs dispensable) is counterintuitive in network science because most work has identified network hubs as important. The highly connected genes in our common genetic architecture drive the insight that the lossy compression of CRISPR libraries is possible and it helps us understand which measurements we might be able to eliminate instead of the measurements that we want to keep.

While screens of single genes in well-behaved cancer cell lines are easy to perform at the genome-wide scale, many contexts exist where the experimental coverage requirements of genome-wide libraries are logistically challenging, if not impossible. Lossy compression changes the experimental calculus by allowing a subset of a genetic library to predict unmeasured gene phenotypes. The largest caveat in our approach is that it is lossy, and therefore has less information than a genome-wide library. However, lossy compression also enables orders of magnitude reductions in library size, i.e. 200 genes and not 20,000. This is most important to consider in the case of the currently accurate null predictions that we are already making in common nonessential genes. Evidence in yeast and worms suggests that broader environmental and developmental contexts elicit more phenotypes across single gene knockouts^42,43^. Thus, the number of “common nonessentials” decreases as more screening conditions are tested. A completely novel phenotype from a novel screen in a common nonessential gene might be challenging to predict with our current lossy 200 compression set. However, new screens are being completed all the time. At the time of our writing, nearly 1000 different mammalian cell lines have already been screened by the Broad and Sanger institute. Only 1144 contexts were needed to identify a measurable phenotypes in every yeast gene^43^. While the human genome will likely require more contexts to saturate phenotypic predictions, lossy sets can be rederived in real time to capture new biology and extend our lossy approach as new data arises. Despite this, many common nonessential genes belong to paralogous gene families and contain genes that are likely to buffer each other’s effects upon single sgRNA knockout. Thus, many current common non-essential genes may never have a measurable phenotype with a single sgRNA^33^ and our current lossy 200 approach would be sufficient to predict phenotypes in these genes in any context. This buffering provides a biologically plausible bound to potential false negatives in lossy compression predictions of common nonessentials.

Despite these minor but important caveats, lossy compression has enormous potential. Chemogenomic screens that kill cells can bottleneck libraries by 10-fold at a 90% inhibition of cell viability. These bottlenecks in screens can raise coverage requirements for genome scale screens to as many as 1 billion cells, a challenging bar. Beyond gene-drug interaction screens, pairwise genetic epistasis maps require *n^2^/2* unique constructs^18^. To make a genome-wide pairwise interaction map (even ignoring the challenge of cloning a pairwise library of 1e8 gene pairs), coverage requirements of 3 sgRNAs per gene, 500 cells/sgRNA and 500 reads/sgRNA create an impossibly large screening challenge that requires a population greater than 10^10^ cells. Because thousands of unmeasured phenotypes can be predicted with hundreds of CRISPR measurements, lossy compression sets are positioned to expand the experimental landscape of possibilities in mammalian functional genomics by enabling genome-scale experiments that were previously impossible.

## Supplemental Information

**S1-a Figure S1.**
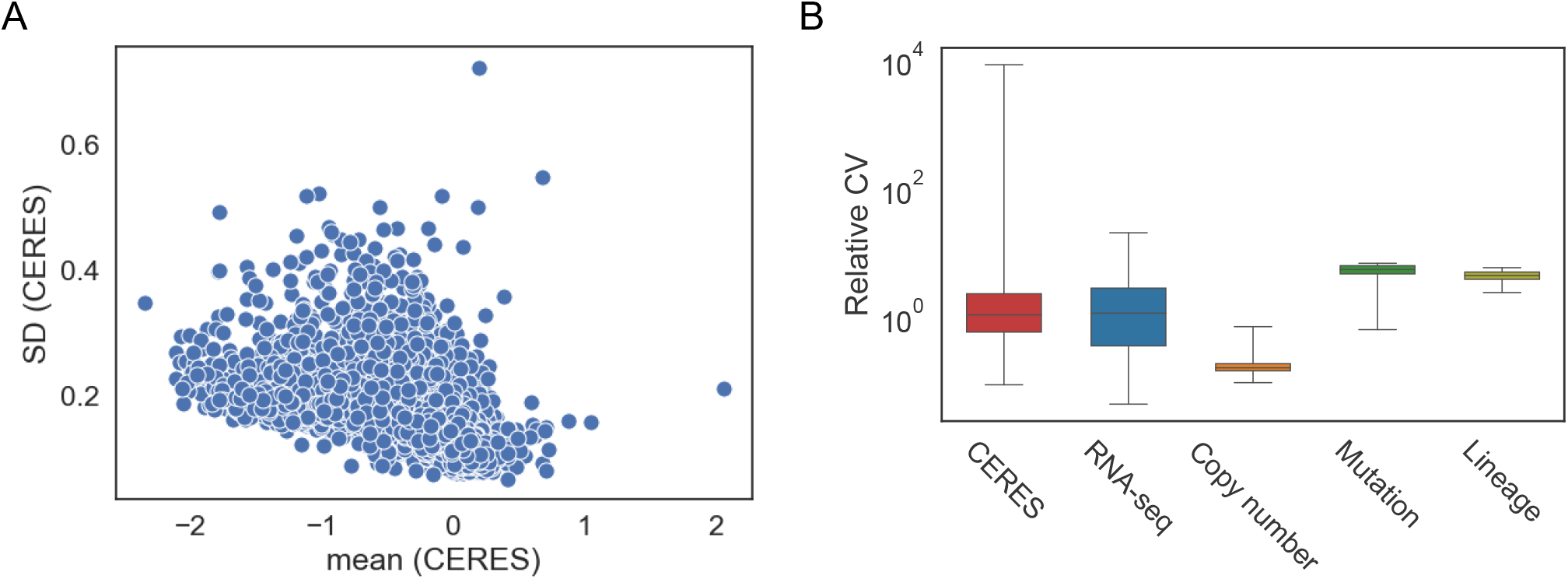
Baseline statistics of DepMap datasets. (A) Scatter plot of standard deviation (SD) versus mean of CERES scores. (B) Relative coefficient of variation (CV) of each data source. Relative CV is calculated as the ratio of standard deviation to the absolute value of the mean.

**S1-b Figure S2.**
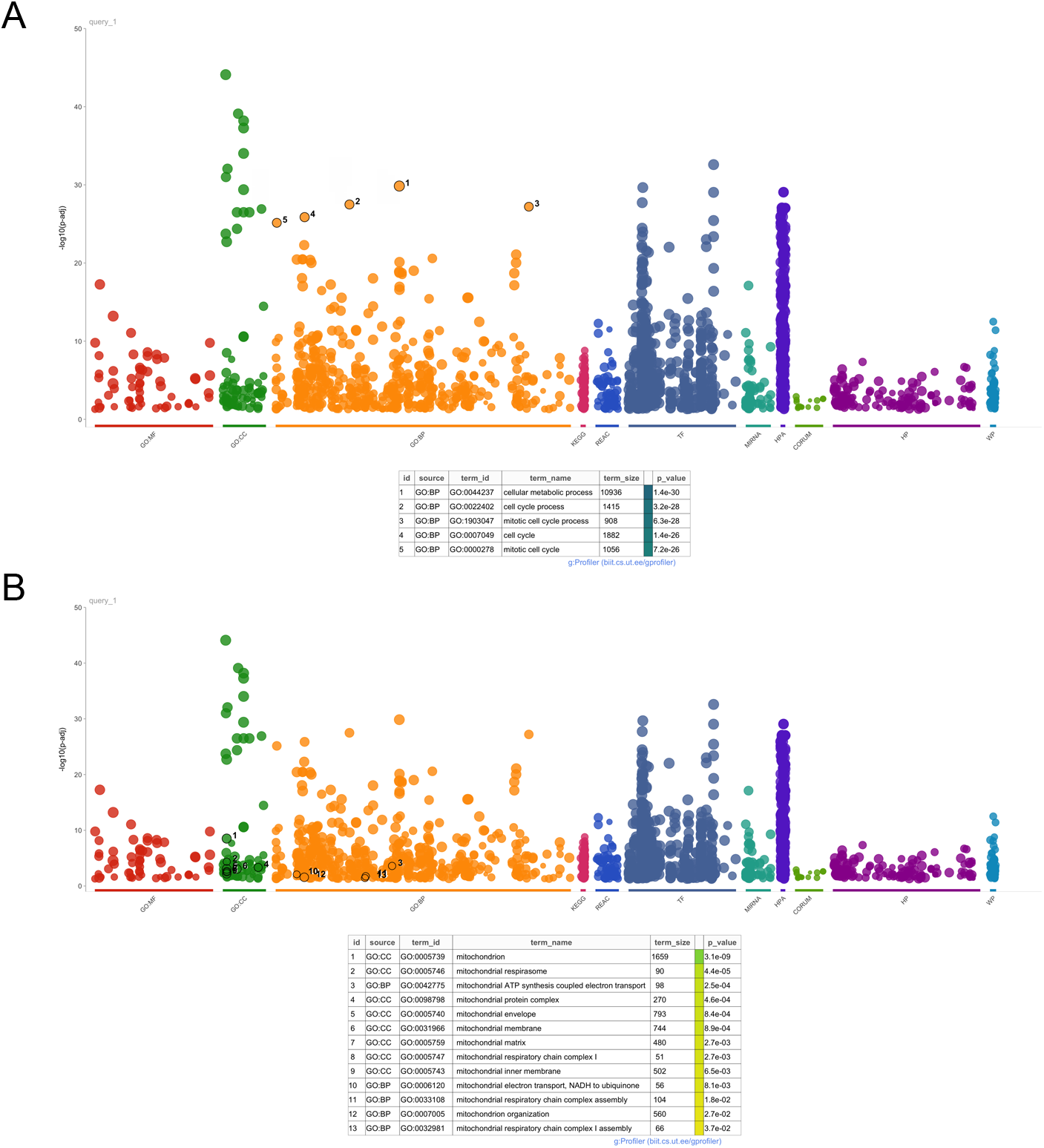
Gene set enrichment of input datasets. Gene set enrichment of gene ontology terms related to biological process (A) and mitochondrial terms (B) of target genes.

**S1-c Figure S3.**
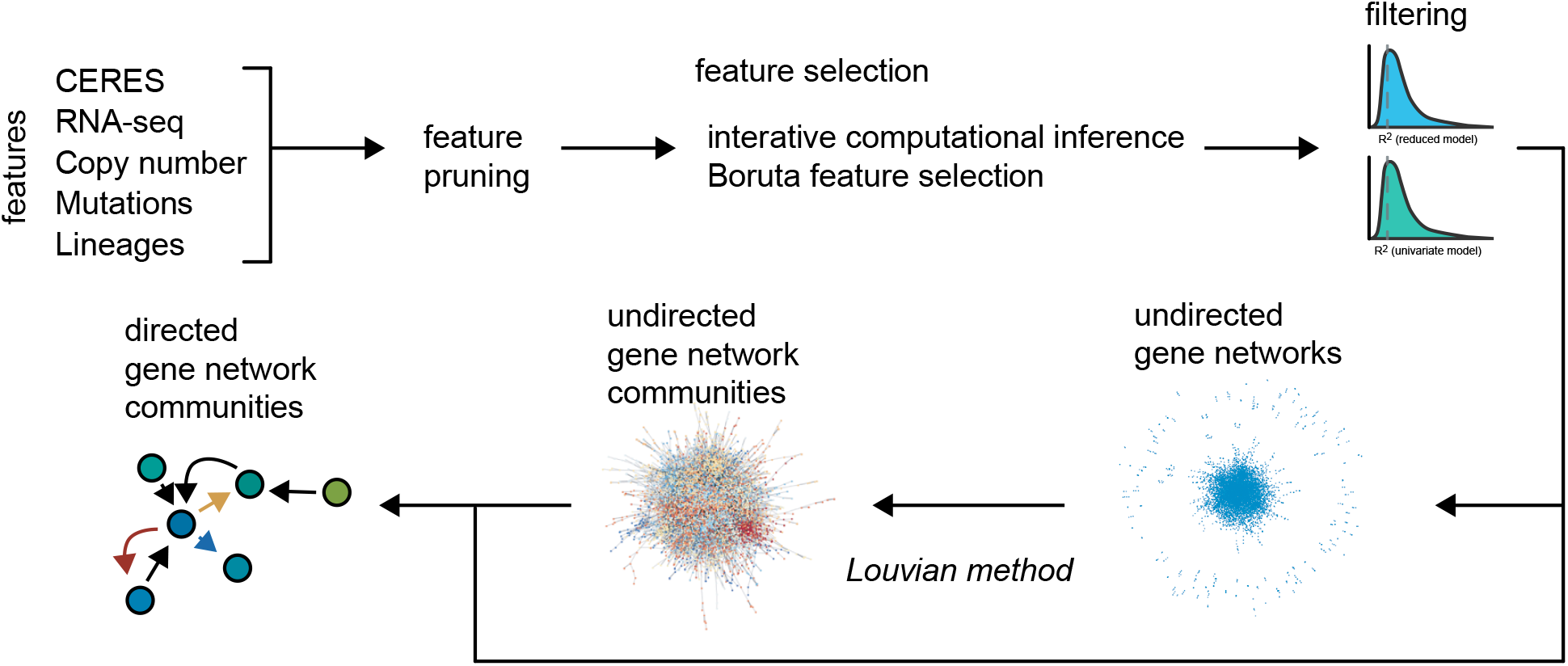
Schematic of model and network building. Data sources (CERES scores, RNA-seq, copy number, mutations, and lineage) are used as features. Prior to model building, invariant/low variant features and non-expressed genes are pruned. The features are selected via an iterative processed followed by Boruta feature selection. Models are fit to the final reduced set of features and further filtered based on model scores. The feature-target genes were aggregated into an undirected gene network, of which communities are identified using the Louvian method. The communities are defined as directed networks based on the feature-target gene relationships. See Methods for more details.

**S1-d Figure S4.**
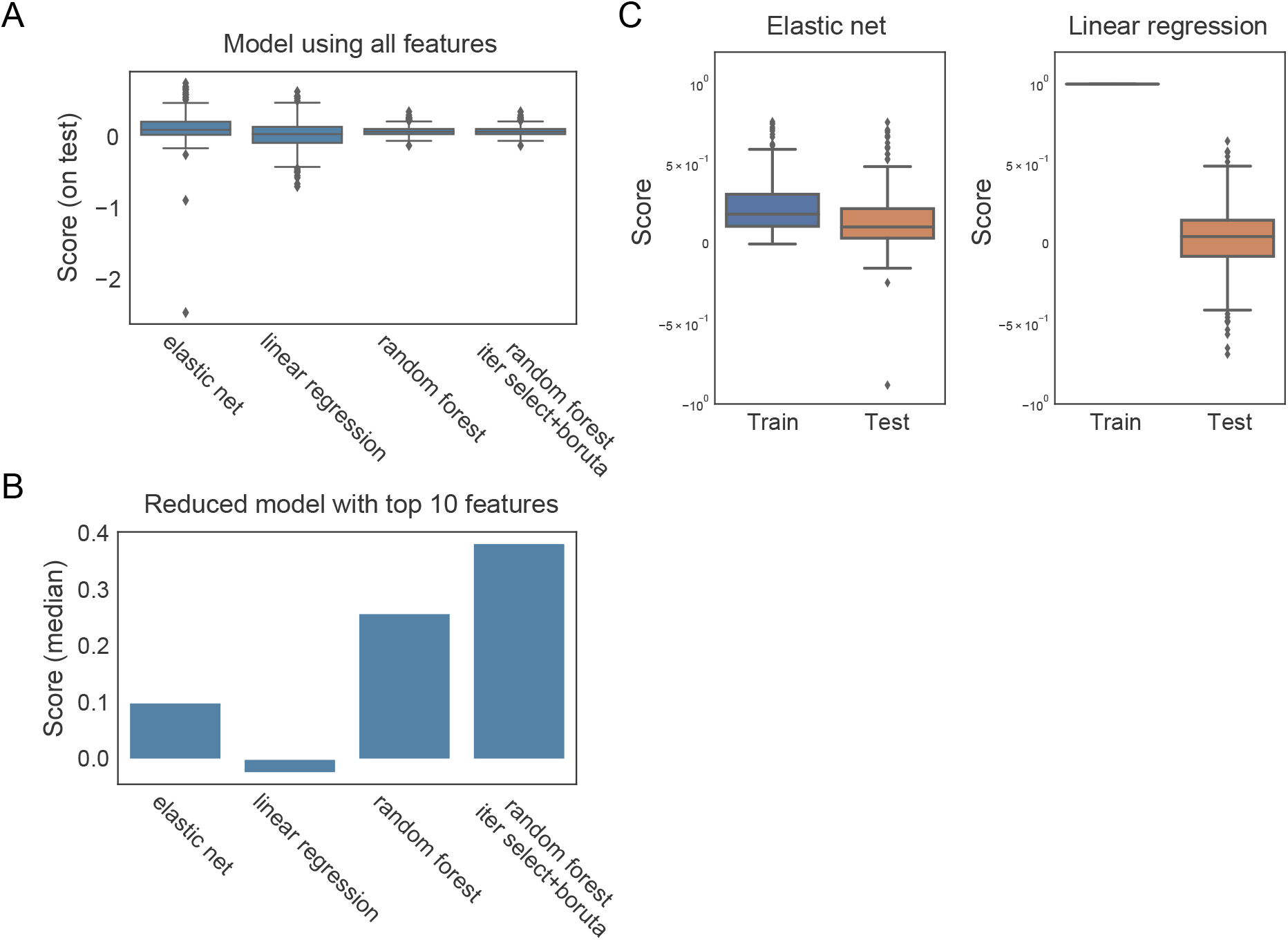
Comparison in performance across different machine learning algorithms. (A) Comparison of model score on test set using all features based on different ML algorithms: elastic net, linear regression, random forest, or random forest with iterative/Boruta selection. (B) Comparison of model scores across different ML algorithms built based on the top 10 features. (C) Model score for train and test set for elastic net (left) and linear regression (right) models based on all features.

**S1-e Figure S5.**
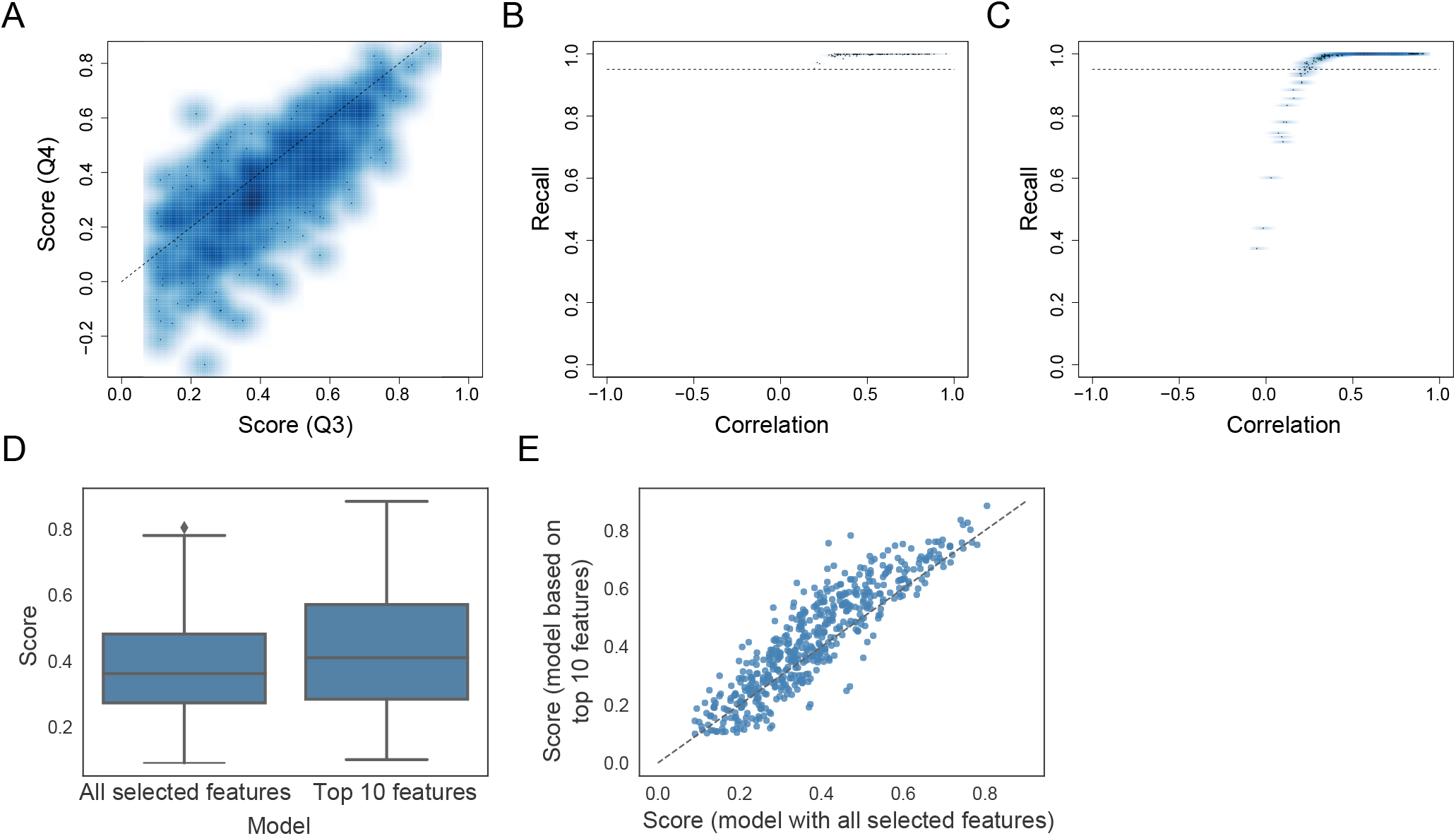
Validation of model performance. (A) Scatter plot of model scores on the 19Q4 vs 19Q3 test sets. Models are trained on the 19Q3 training set. (B-C) Recall versus correlation for 19Q3 (B) and 19Q4 (C) test sets. (D-E) Top 10 features are sufficient and shows comparable model performance to models with all significant features, as shown by box-whisker plot (D) and scatter plot (E).

**S2-a Figure S6.**
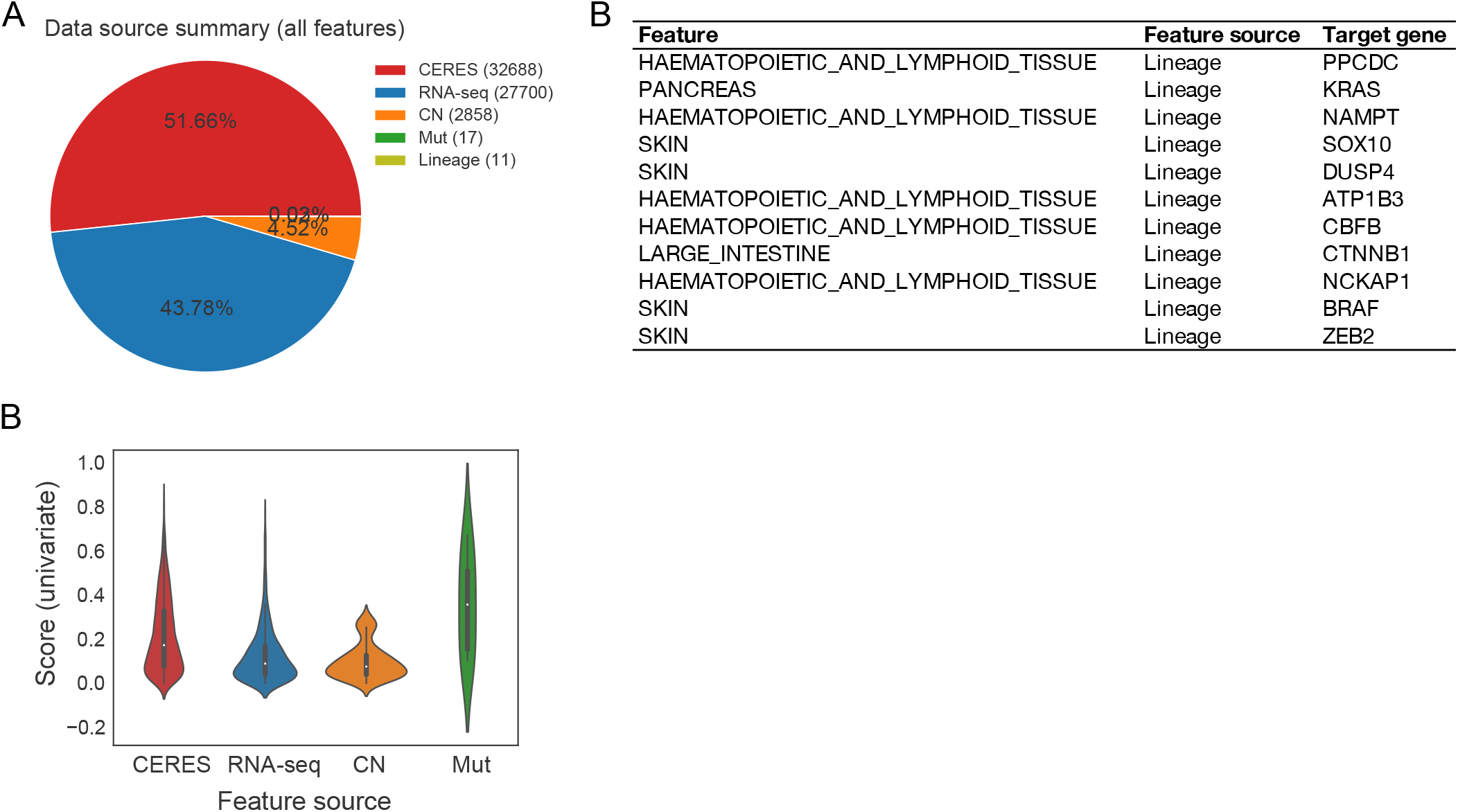
Model feature summaries. (A) Classification of all features of all models by data source. (B) Significant lineage features and their target gene. (C) Univariate model scores as violin plots grouped by the data source of features.

**S2-b Figure S7.**
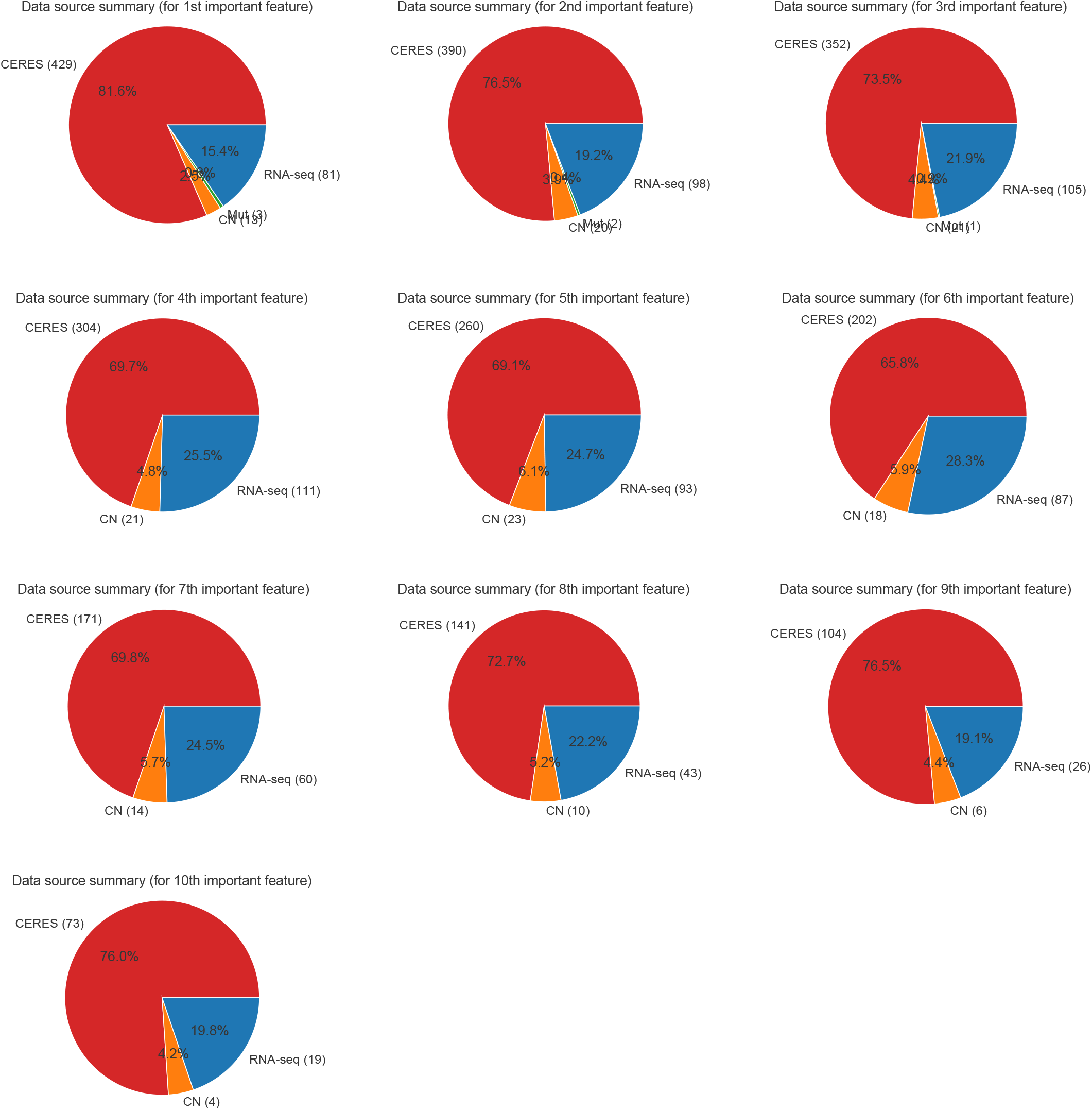
Contribution of data sources for top 10 features. Pie chart showing breakdown of the data source type of features contributing to model prediction, for each *n*th most important feature.

**S3-a Figure S8.**
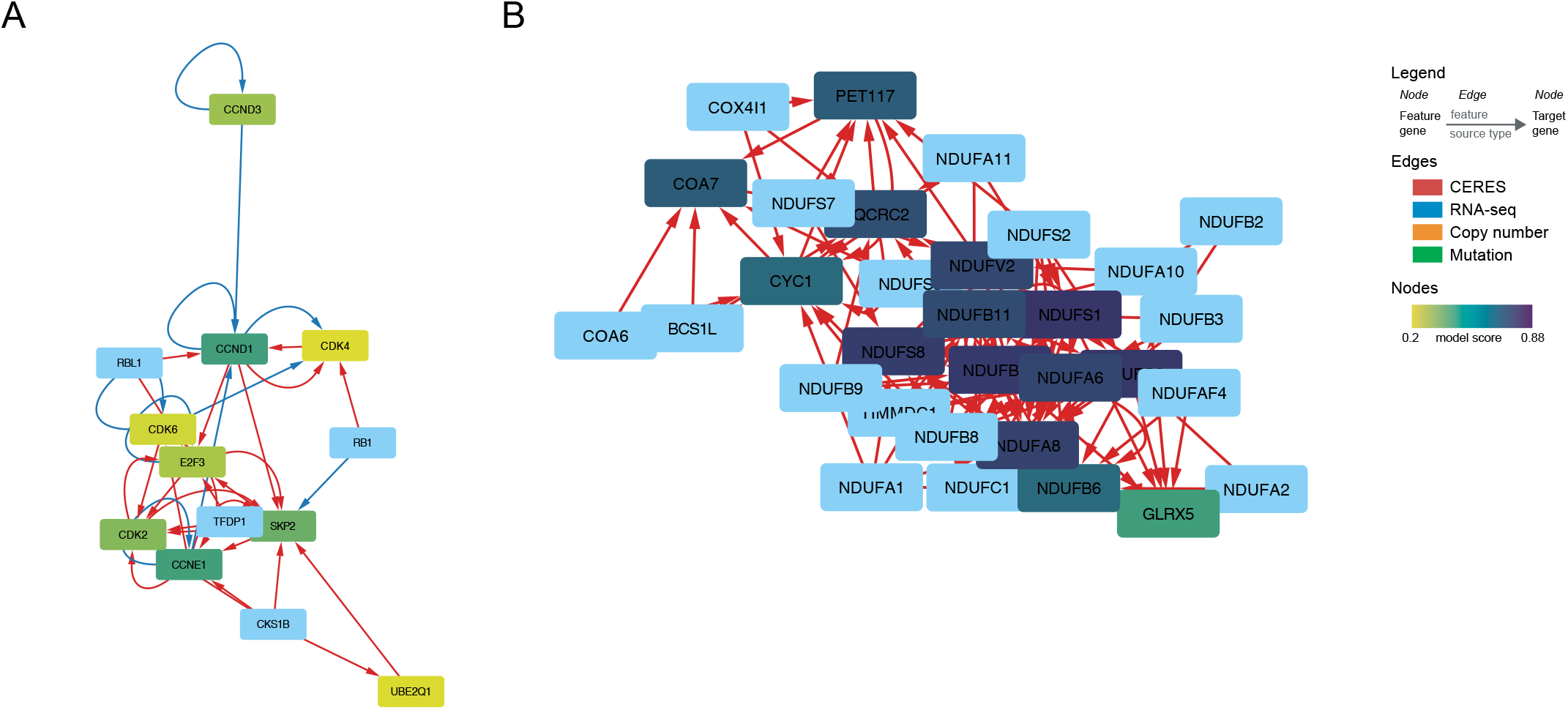
Selected network communities. CDK4 (A) and mitochondrial (B) networks extracted from model/network building. Nodes denote genes and edges denote feature-target gene relationships. Node colors are based on the score of the top 10 feature model of the corresponding gene as target.

**S4-a Figure S9.**
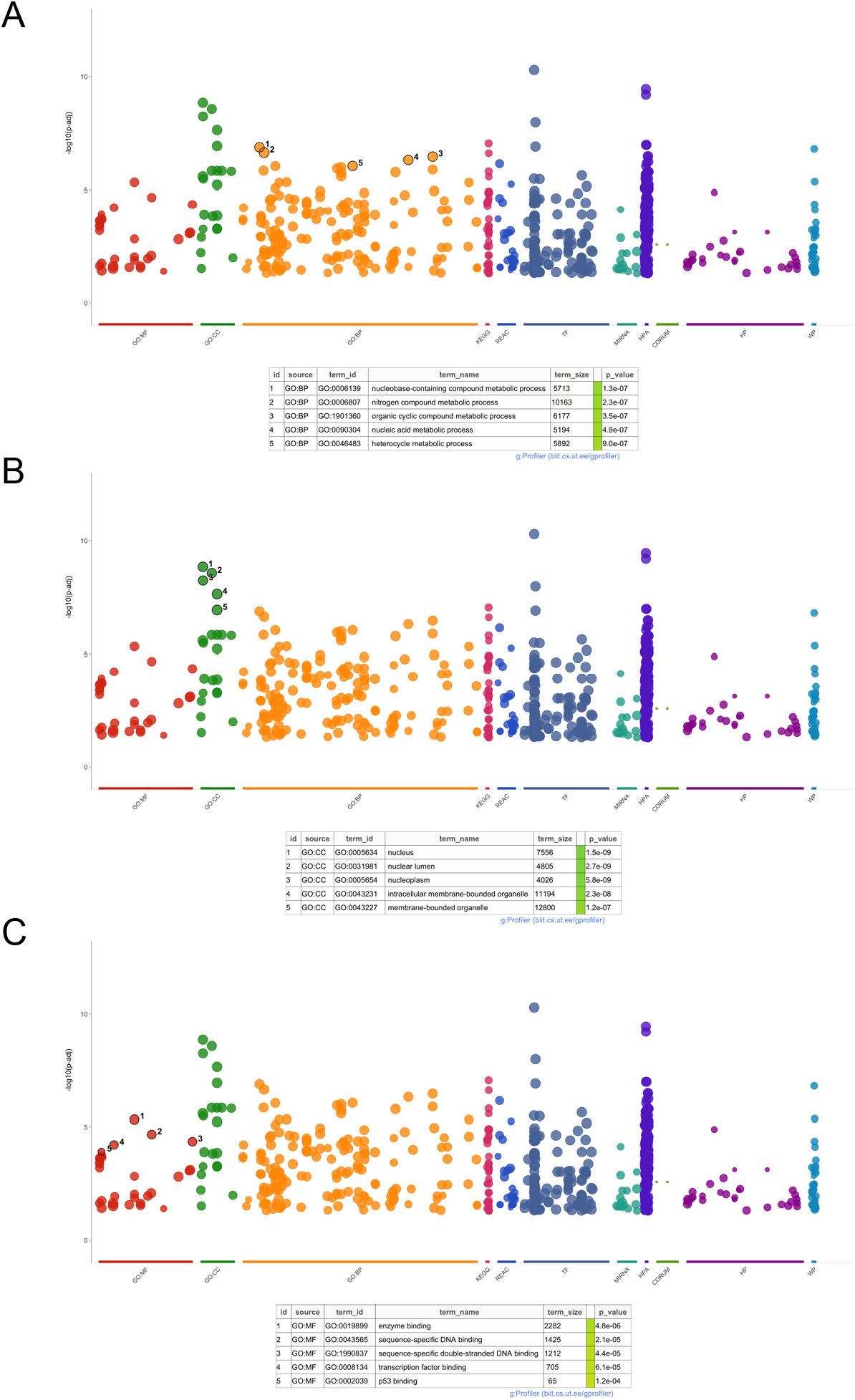
Gene set enrichment of L200 lossy gene sets. Gene set enrichment of gene ontology terms related to biological process (A), molecular function (B), and cellular compartment (C) of L200 lossy gene sets, using gProfiler.

**S4-b Figure S10.**
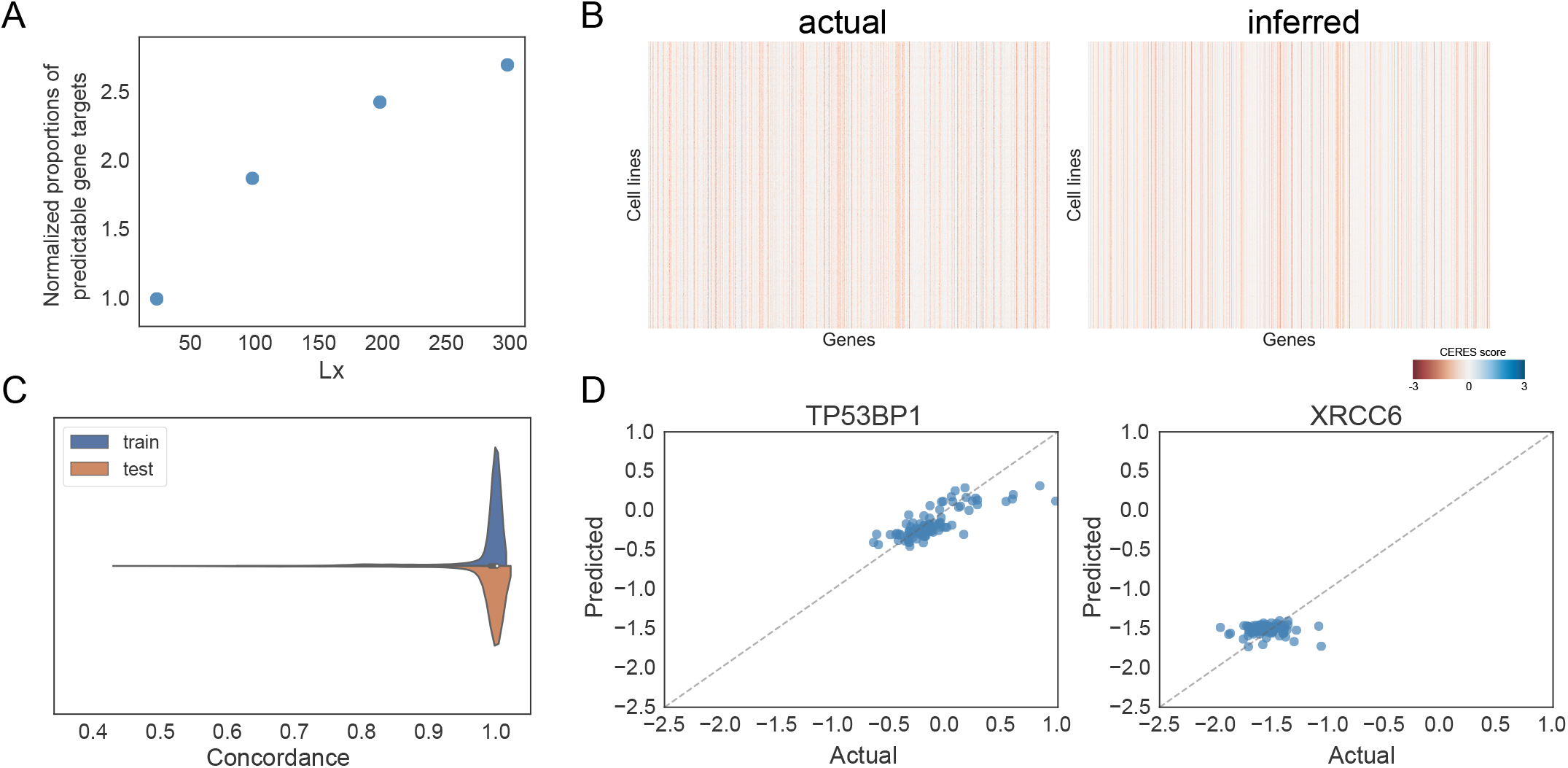
Validation of lossy compression. (A) Saturation analyses based on lossy L25, 100, 200, and 300 lossy gene sets. Y-axis shows the normalized proportion of predictable gene targets, where predictable is defined as target genes with recall greater than 0.95. (B) Genome-wide comparison between actual and inferred CERES scores on the training set. (C) Concordance between actual and predicted CERES scores for train and test sets. (D) Scatter plot of predicted versus actual for selected genes *TP53BP1* and *XRCC6*.

## Notes

https://github.com/pritchardlabatpsu/cnp_dev

### Competing Interest Statement

The authors have declared no competing interest.

